# Structural basis of Ebola virus nucleocapsid assembly and functions

**DOI:** 10.1101/2024.02.19.580633

**Authors:** Yoko Fujita-Fujiharu, Shangfan Hu, Ai Hirabayashi, Yuki Takamatsu, Yen Ni Ng, Kazuya Houri, Yukiko Muramoto, Masahiro Nakano, Yukihiko Sugita, Takeshi Noda

## Abstract

The Ebola virus, a member of the *Filoviridae* family, causes severe hemorrhagic fever in humans. Filamentous virions contain a helical nucleocapsid responsible for genome transcription, replication, and packaging into progeny virions. The nucleocapsid consists of a helical nucleoprotein (NP)–viral genomic RNA complex forming the core structure, to which VP24 and VP35 bind externally. Two NPs, each paired with a VP24 molecule, constitute a repeating unit. However, the detailed nucleocapsid structure remains unclear. Here, we determined the nucleocapsid-like structure within virus-like particles at 4.6 Å resolution using single-particle cryo-electron microscopy. Mutational analysis identified specific interactions between the two NPs and two VP24s and demonstrated that each of the two VP24s in different orientations distinctively regulates nucleocapsid assembly, viral RNA synthesis, and intracellular transport of the nucleocapsid. Our findings highlight the sophisticated mechanisms underlying the assembly and functional regulation of the nucleocapsid and provide insights into antiviral development.

## Introduction

The Ebola virus (EBOV), which belongs to the *Filoviridae* family in the *Mononegavirales* order, causes fatal hemorrhagic fever in both humans and non-human primates^1^. Although recent advancements in the availability of Food and Drug Administration-approved vaccines and antibody drugs exist^2^, their therapeutic targets are limited to glycoproteins (GP), which are responsible for virus entry into target cells. Therefore, exploring the structures and functions of other viral proteins and understanding the molecular mechanisms underlying EBOV replication are required for the development of novel antiviral therapeutics.

The EBOV nucleocapsid, responsible for the transcription and replication of non-segmented, single-stranded, negative-sense viral genomic RNA (vRNA), is composed of a nucleoprotein (NP), VP24, VP30, VP35, and polymerase L, in which a helical NP–vRNA complex serves as the core structure^1^. VP24 and VP35, essential for the formation of nucleocapsid-like structures (NCLSs)^3,4^, bind externally to the helical NP–vRNA helix^5,6^, forming protrusions on its surface^7–10^. VP35, which is analogous to the P protein of other mononegaviruses^11^, is a polymerase cofactor that bridges the NP–vRNA helix and L during vRNA synthesis^12–15^. VP35 also binds to nascent NPs to prevent oligomerization before forming the helical NP–vRNA complex^16,17^. In contrast, the characteristics of VP24 remain poorly understood owing to the absence of analogs in mononegaviruses other than filoviruses. Currently, it is established that VP24 suppresses transcription and replication of the vRNA^18,19^. Additionally, it is essential for the intracellular transport of nucleocapsids and, thus, genome packaging into progeny virion^20,21^. This suggest that VP24 may switch nucleocapsid functions from RNA synthesis to vRNA transport for virion formation. However, the underlying structural basis for its regulatory functions remains elusive because of the lack of knowledge regarding the detailed nucleocapsid structure.

Previously, we determined the high-resolution structure of the EBOV NP–RNA complex at 3.6 Å resolution using single-particle cryo-electron microscopy (cryo-EM) and identified essential interactions on NP for the formation of the helical NP–RNA complex and for the transcription and replication of the vRNA^22,23^. In addition, studies using cryo-electron tomography (cryo-ET) determined the nucleocapsid structure in authentic EBOV particles and the NCLS structure in virus-like particles (VLPs) at 9.1 Å and 7.3 Å resolution, respectively^8^. These studies generated a model of the nucleocapsid structure, in which a repeating unit consists of two NPs with two protruding VP24s in different orientations^8^. However, the absence of a high-resolution structure of the nucleocapsid hinders our understanding of the structural characteristics and functions of the EBOV nucleocapsid assembly.

The aim of this study was to report the *in situ* structure of EBOV NCLS in VLPs at the 4.6 Å resolution. These results advance our understanding of the molecular mechanisms underlying EBOV replication and contribute to the development of novel antiviral therapeutics.

## Results

### Cryo-EM structure of the EBOV NCLS in VLPs

To better understand the molecular interactions among nucleocapsid components that regulate the assembly and function of the EBOV nucleocapsid, we attempted to determine the higher-resolution structure of the EBOV NCLS within VLPs using single-particle cryo-EM, which differs from previous studies using cryo-ET^7,8^. Consequently, we co-expressed NP, VP24, VP35, VP40, and GP in a human cell line and produced NCLS-containing VLPs that were morphologically indistinguishable from authentic EBOV particles^24^. Initially, we employed a single-particle analysis, imposing helical symmetry, and determined the helical NCLS structure inside the VLP at an overall resolution of 7.5 Å (Fig. 1A, Extended Data Fig. 1). Our cryo-EM map shows a good fit with the cryo-ET maps of the NCLS in the VLP and the nucleocapsid in authentic virion^8^ (Fig. 1B). It maintains a consistent helical parameter (Extended Data Table 1), suggesting that our cryo-EM structure accurately represents the native conformation of the EBOV nucleocapsids. The structure of the NP–RNA complex located within the inner layer of the NCLS is analogous to the high-resolution structure we reported previously^22^ (Extended Data Fig. 2). The external protrusions on the helical NP–RNA complex were presumed to be two VP24 molecules, as demonstrated previously^8^. We also detected additional map densities outside the VP24 protrusions that remained uncharacterized, as reported in cryo-ET analysis^8^ (Fig. 1C, shown in dark gray). Next, to identify the molecular interactions within the nucleocapsid, we focused on its repeating unit, composed of two NPs and two VP24s. We achieved a higher-resolution structure by expanding the helical symmetry and further processing the repeating unit (Extended Data Fig. 1). The resulting map, with a 4.6 Å resolution, facilitated the identification of the NP–RNA complex that constitutes the inner core of the nucleocapsid and the positioning of VP24s on the NP–RNA complex (Fig. 1C,D). Unfortunately, we could not identify the outer map of the VP24 protrusions owing to the limited resolution (Fig. 1B, shown in dark gray). The repeating unit of the NCLS was composed of two adjacent copies of NP-associating RNA and two differently oriented VP24 molecules (Fig. 1C,D): NP-1 (shown in pink) featured a large protrusion of additional density on the outside of the core, which included one copy of VP24-1 (shown in lime green), and NP-2 (shown in beige) had a smaller protrusion that contained one copy of VP24-2 (shown in violet).

**Fig. 1.**
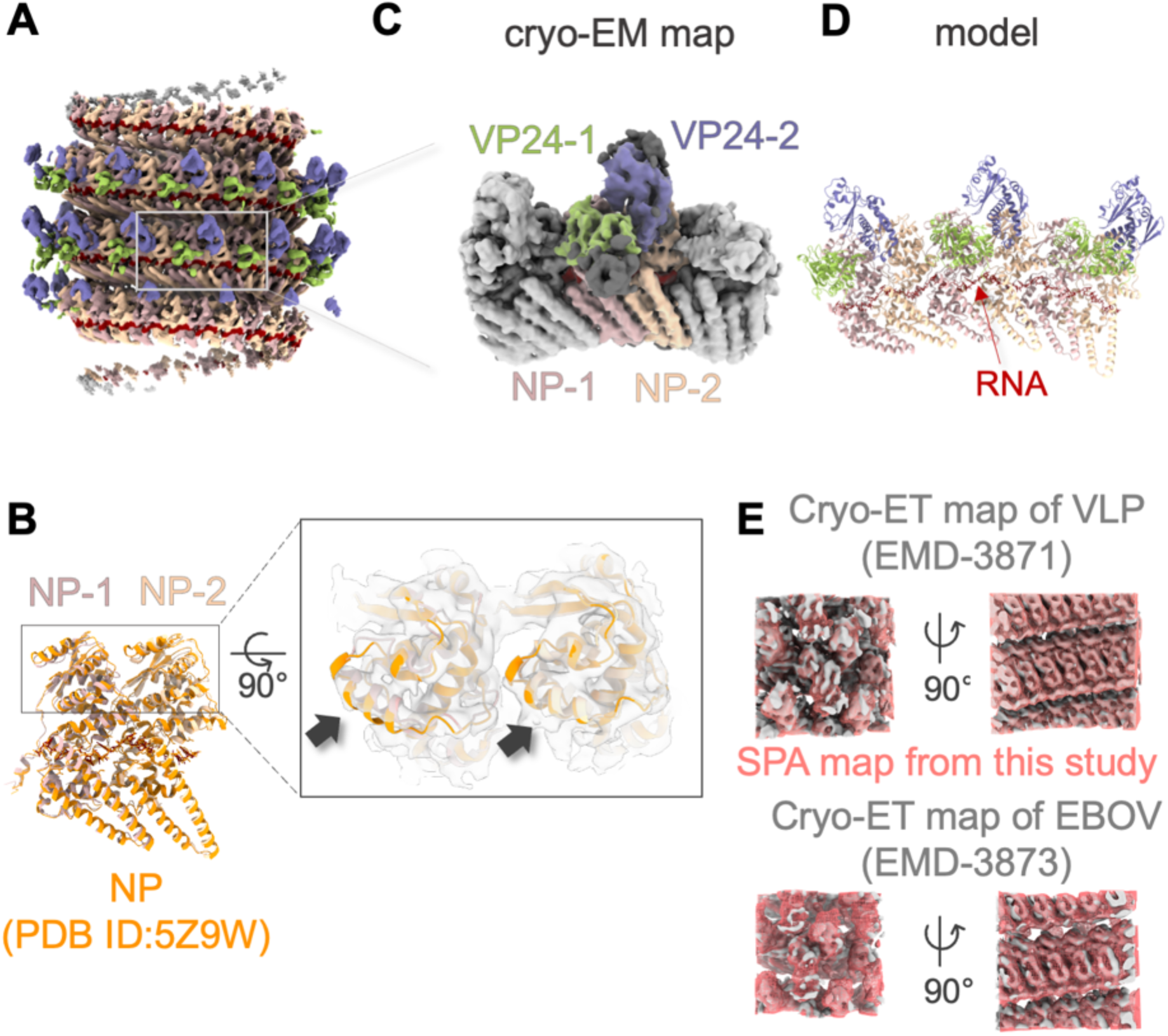
Cryo-EM structure of EBOV nucleocapsid-like structure (NCLS) in VLPs. A. Helical reconstruction of the NCLS complex at 7.5 Å resolution, with RNA colored in red, nucleoproteins (NP-1 and NP-2) in pink and beige, and viral proteins VP24-1 and VP24-2 in lime green and violet. B. Comparison of the NP structures derived from this study with the previously published purified NP structure (PDB ID: 5Z9W), shown in orange. The left image shows a side view, while the inset displays a 90° rotated top view with the cryo-EM map from this study. The arrow indicates a local structural difference between these NP structures. C. Symmetry-expanded and focus-refined cryo-EM map at 4.6 Å resolution showing the arrangement of NPs and VP24 molecules, with VP24-1 in lime green and VP24-2 in violet alongside NPs (NP-1 in pink and NP-2 in beige). Adjacent repeating units are shown in light grey; areas, where the model could not be assigned, are shown in dark grey. D. Atomic model of the NCLS. The color scheme for NPs and VP24s matches that of Fig. 1A. E. Overlay of the cryo-electron tomography (cryo-ET) map of the VLP (EMD-3871) and the authentic virus (EMD-3873) compared with the cryo-EM map obtained in this study, showing structural congruence.

The overall architecture of the NP core closely resembled that of the purified NP–RNA complex (Fig. 1E). The root-mean-square deviation (RMSD) of backbone atoms was 1.05 Å between NP-1 and the purified NP (Protein Data Bank (PDB) ID: 5Z9W)^22^, and 1.05 Å between NP-2 and the purified NP. Notably, a marginal difference was observed in the local structure of the N-terminal lobe domains of both NP-1 and NP-2 (Fig. 1B indicated by arrows). This observation suggested that the NPs adjust their position at the inter-rung interface after binding to VP24 and VP35, as indicated in a cryo-ET study^8^.

### NP–VP24 interaction in the NCLS

The NP–VP24 interaction involves three distinct interfaces. One interface is formed via the interaction between the C-terminal helix of NP-1 and a loop on VP24-1 (Fig. 2A,B), whereas the other two interfaces primarily rely on electrostatic interactions between the N-terminal lobes of NPs and VP24s (Fig. 2C-E). The 169-173 loop region of VP24, in which the YxxL motif is included, is recognized as a crucial site for NP–VP24 interactions and intracellular transport of nucleocapsids^21,25^. Our structure revealed an interaction between the C-terminal α-helix 18 of NP-1 and the 169-173 loop of VP24-1 (Fig. 2B). Notably, the loop region exhibited considerable flexibility, and the local resolution of this region was limited (Extended Data Fig. 3A,B); hence, the precise positioning of each amino acid in this loop could not be determined.

**Fig. 2.**
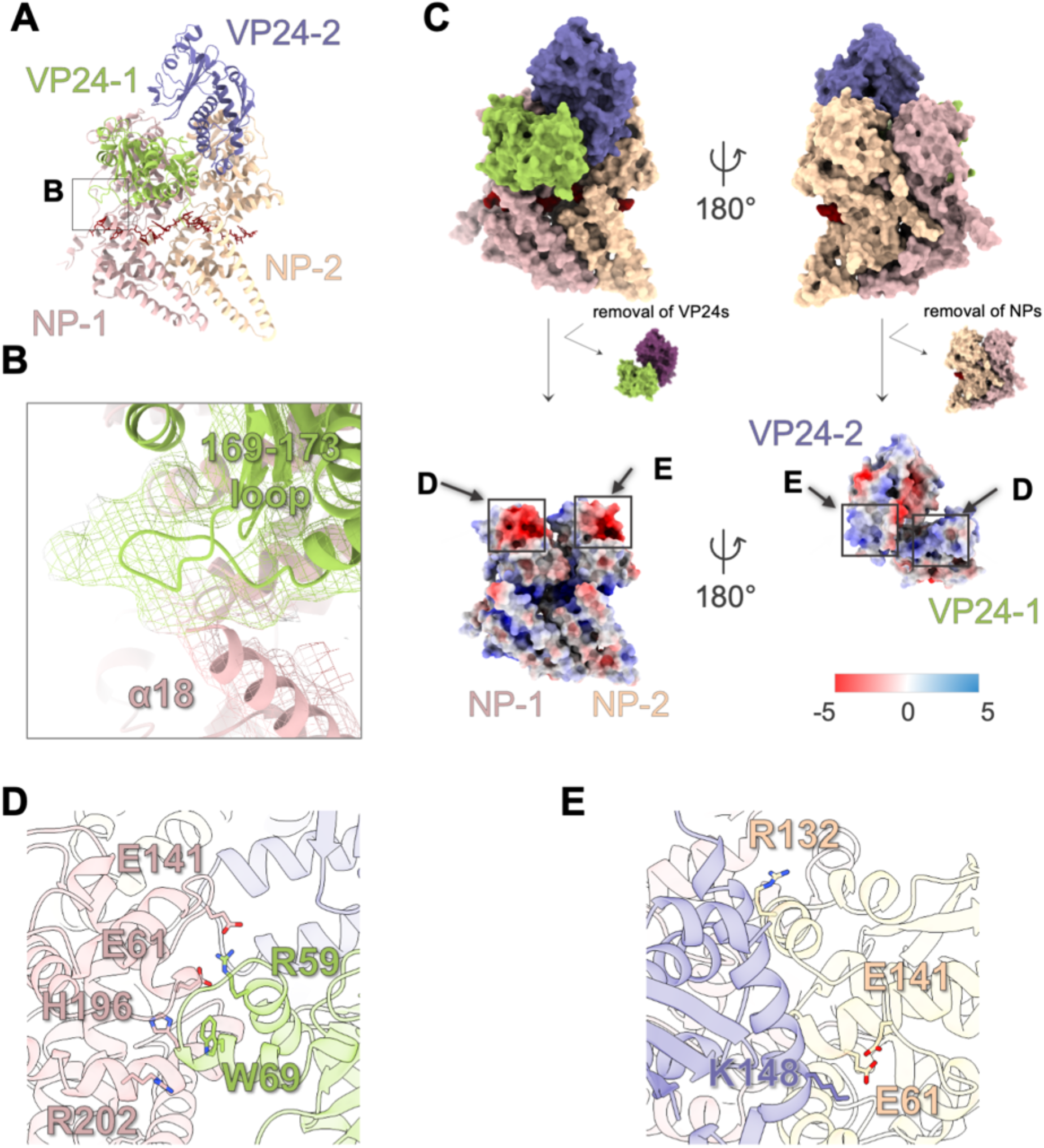
Detailed interactions between NPs and VP24s within the Ebola virus nucleocapsid-like structure. A. Atomic model of the NCLS showing the helical repeating unit containing two NPs (NP-1 in pink and NP-2 in beige) and two VP24 molecules (VP24-1 in lime green and VP24-2 in violet). B. Close-up view of the interface between the 169-173 loop of VP24-1 and the α18 helix of NP-1, with the mesh representing the cryo-EM map. C. Electrostatic surface potential maps of the NCLS from two orientations, 180° apart. The maps show the distribution of charges around NP-1 and NP-2, with VP24-1 and VP24-2, respectively. Electrostatic surface potential was calculated using the Delphi web-server^39^. Scale from –5 (red) to +5 kcal mol^−1^ e^−1^ (blue). D. Close-up view of the NP-1 and VP24-1 interface, detailing electrostatic contacts. Residues involved in potential salt bridge and pi-cation interactions labeled and presented in stick representation. E. Close-up view of the NP-2 and VP24-2 interface.

Examining the other interfaces mediated via the N-terminal lobe of the NPs and each VP24 subunit, electrostatic interactions emerge as the predominant driving force for their binding (Fig. 2C). In our structural model, R59 of VP24-1 is inserted into the negatively charged region of NP-1, which is composed of residues E61 and E141, forming a possible salt bridge (Fig. 2D). Additionally, H196 of NP-1 is located near W69 of VP24-1, suggesting a potential interaction involving pi-cation stacking. NP-1 R202 is positioned in a ‘clipping’ VP24-1 manner. Examining the adjacent pair of NP-2–VP24-2, it is observed that R132 of NP-2 is located where it could interact with the negatively charged region of VP24-2 (Fig. 2C,2E). Notably, K148 of VP24-2 is inserted into the negatively charged region comprising residues E61 and E141 of NP-2, resembling the positioning of R59 in the adjacent VP24-1. This observation suggested that the negatively charged surface of the NP N-terminal lobe is important for binding both VP24s.

### VP24–VP24 interactions in the NCLS

Compared to our two VP24 models, a previously reported crystal structure of the EBOV VP24 dimer^26^ showed substantial differences in arrangement, potentially owing to its crystal packing (Extended Data Fig. 3C). In addition, the crystal structure shows different conformations with RMSDs of 3.66 Å for VP24-1 and 2.35 Å for VP24-2 (Extended Data Fig. 3C). Thus, the VP24 dimer in the crystal packing does not represent the arrangement and conformation within the NCLS; however, it may instead depict an interferon antagonist form^27^. Notably, the two VP24s observed in our NCLS structure exhibit a substantial difference in their conformation at the main chain level, potentially owing to their association with each NP, with RMSD of 1.37 Å (Extended Data Fig. 3D).

In our model, no apparent inter-unit interaction, such as that between VP24-1 in a repeating unit and VP24-2 in a neighboring repeating unit, is observed. This is because of the distinct positioning of each VP24 unit, separated from those in neighboring units. Notably, an atomic model could not be constructed for the N– and C-termini (1-9 a.a. and 232-251 a.a. for VP24-1, and 1-27 a.a. and 232-251 a.a. for VP24-2) (Extended Data Fig. 3A,E), leaving the possibility of interactions via this region open. In contrast, there seems to be an intra-unit interaction between VP24-1 and VP24-2 (Fig. 3A), with a specific region showing a potential contact based on a cut-off distance (>7 Å). Within this interface, VP24-2 forms a hydrophobic pocket, in which a hydrophobic residue, I35 of VP24-1, is inserted (Fig. 3B,C). Given that VP24-VP24 interactions are limited to this hydrophobic interaction with I35E (Fig. 3A), the binding affinity between the two VP24s within the NCLS would be relatively weak.

**Fig. 3.**
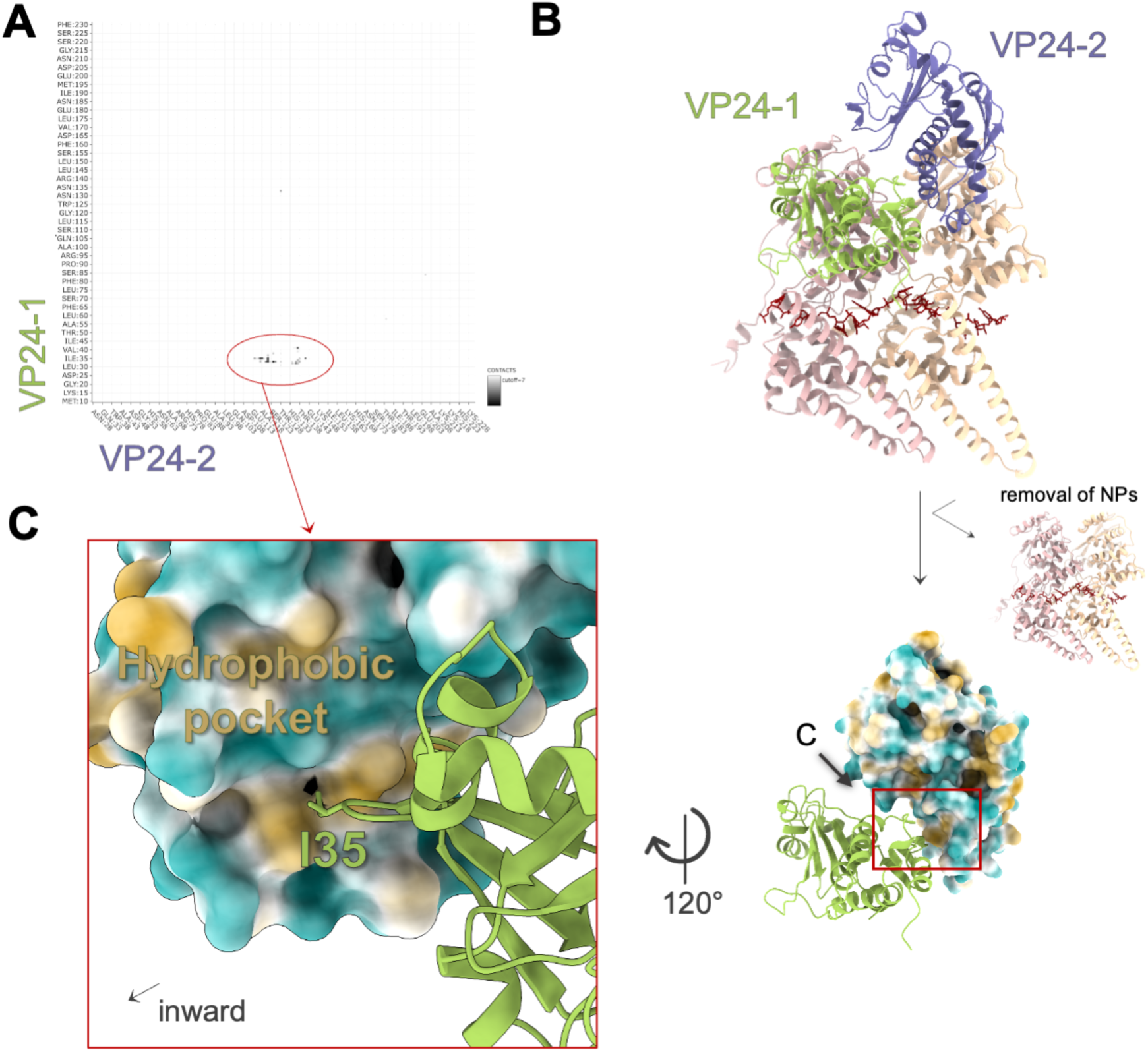
Interactions between VP24s within the NCLS. A. A contact map showing the interaction sites between VP24-1 and VP24-2. The red circle indicates a key area where contacts are enriched. B. The atomic model showing the spatial arrangement of VP24-1 (lime green) and VP24-2 (violet) in ribbon representation, with the molecular lipophilicity potential map of the VP24-2 structure (in surface representation) colored in blue (hydrophilic) to orange (hydrophobic). C. Close-up view of the hydrophobic pocket formed at the interface between VP24-1 and VP24-2, corresponding to the red circled area in the contact map (A). This close-up demonstrates residue I35 of VP24-1 resides within the hydrophobic pocket of VP24-2.

### NP–VP24 interaction required for VP24 recruitment into the inclusion body

To identify the specific residues responsible for NCLS formation and function, we generated eight NP and VP24 mutants based on our structural model. We designed four NP mutants at the interface with VP24-1 (E61K, H196A, and R202A) and VP24-2 (E61K and R132A), three VP24 mutants at the interface with NP-1 (R59A and N171A) and NP-2 (K148A), and one VP24 mutant at the hydrophobic interface with another VP24 (I35E). The E61K of the NP is located within both interfaces. Initially, to investigate the interactions between NP and VP24, we performed an immunoprecipitation (IP) assay using these mutants (Fig. 4A). Among all mutants, the VP24 N171A mutant lost its ability to interact with NP, consistent with previous reports^21^. In contrast, the other mutants exhibited interactions between NP and VP24. This result suggested that the interaction of the 169-173 loop of VP24-1 with the C– terminal α-helix 18 of NP-1 is primarily responsible for the interaction between NP and VP24.

**Fig. 4.**
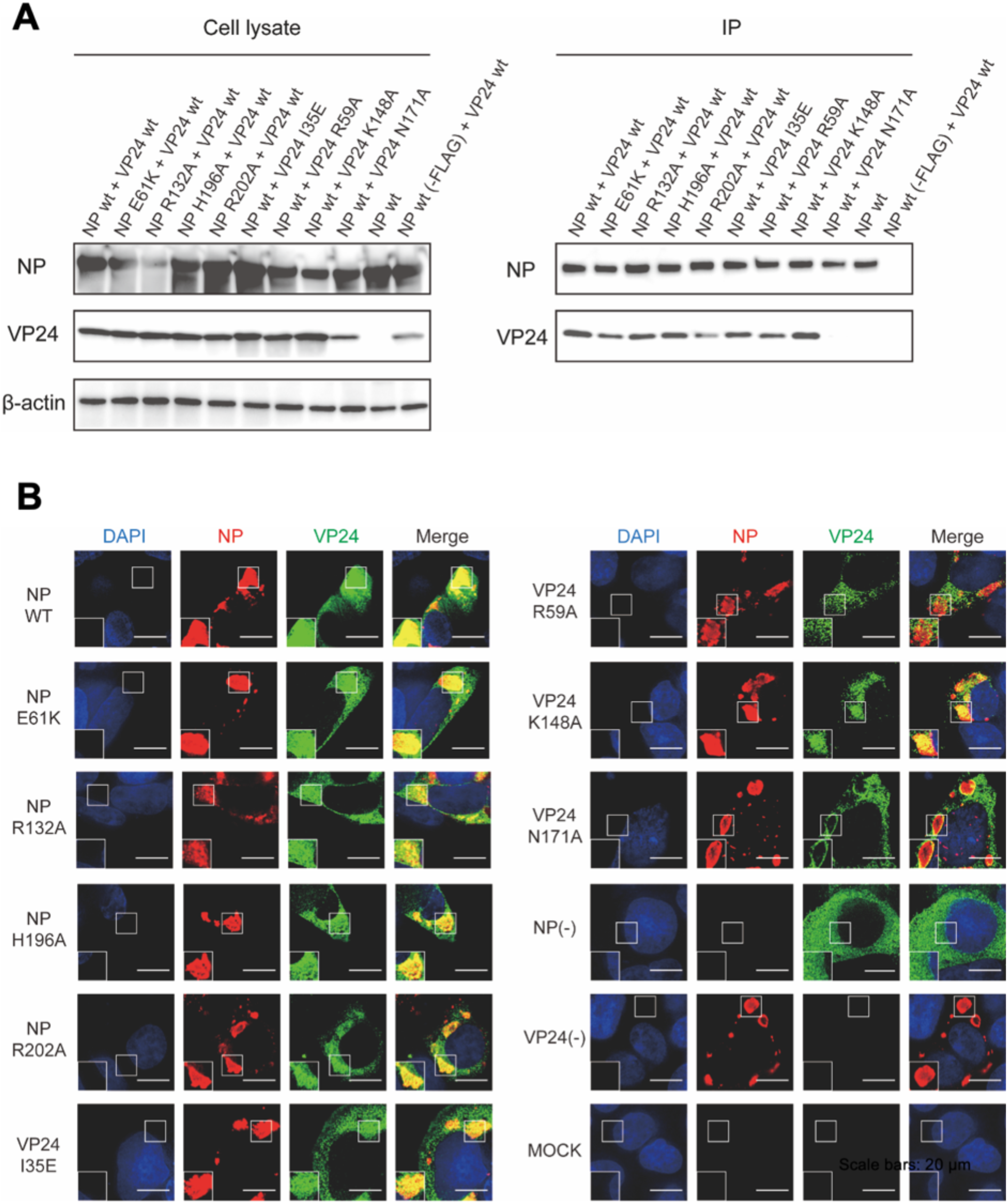
Mutational analysis of NP and VP24 interactions in cells. A. Immunoprecipitation assay using FLAG-tagged NP and VP24 mutants. Left panel; VP24s, co-immunoprecipitated with FLAG-tagged NPs using an anti-FLAG antibody, were analyzed using western blotting using anti-VP24 and anti-FLAG antibodies. Right panel; whole cell lysates were analyzed using western blotting with anti-VP24 and anti-FLAG antibodies. Beta-actin was detected as a loading control. B. Intracellular localization of NP and VP24 mutants in cells co-expressing FLAG-tagged NP, VP24, and VP35.The cells were immunostained using anti-NP (red) and anti-VP24 (green) antibodies. Nuclei were stained with Hoechst 33342 (blue). The absence of NP or VP24 is indicated by (-), and the MOCK control demonstrates cells without plasmid transfection. Scale bars represent 20 µm.

NP expression induces the formation of inclusion bodies (IBs), the sites for viral genome transcription and replication^28^, where VP24 is recruited during virus replication^29^. Therefore, to examine whether these mutants accumulate in IBs, we examined the intracellular localization of the NP and VP24 mutants using an immunofluorescence assay (IFA). When wild-type NP, VP24, and VP35, the minimal components of NCLS formation^3,4^, were co-expressed, VP24 was localized within the IBs formed by NP expression, as reported previously^20,21,29^ (Fig. 4B). VP24 N171A, which showed no interaction with NP as demonstrated by the IP assay, was not localized within IBs. Unexpectedly, VP24 R59A, located at the interface between VP24-1 and NP-1, was not recruited into IBs. All other mutants with mutations at the interfaces between VP24-2 and NP-2 and between VP24-1 and VP24-2, showed the localization of VP24 within the IBs, similar to the wild-type combination (Fig. 4B). These results suggested that VP24 N171 is crucial for the interaction between NP and VP24 and that the interaction between NP-1 and VP24-1 is important for the recruitment of VP24 into IBs.

### Relationship between regulation of polymerase activity and NCLS formation

To investigate the amino acid residues involved in nucleocapsid function, we performed a minigenome assay using the mutants and assessed their impact on the transcription and replication activity of the nucleocapsid. In the presence of wild-type VP24, we observed a significant reduction in the transcription and replication activity, as previously reported^19,30^ (Fig. 5A). The VP24 N171A and R59A mutants, which did not localize within the IBs, showed significantly reduced inhibitory activity compared to VP24 WT, whereas the other VP24 mutants showed similar reductions as observed with wild-type VP24 (Fig. 5A). We were unable to quantitatively evaluate the inhibitory activity of VP24 for most NP mutants because they showed reduced transcription and replication activities, even in the absence of VP24 (Fig. 5B). However, in the NP mutants, the transcription and replication activities of the nucleocapsid composed NP H196A and R202A, which localize at the interface between NP-1 and VP24-1 and show sufficient transcription and replication activity, were mildly inhibited by wild-type VP24 (Fig. 5C). This suggested that the interaction between NP-1 and VP24-1 plays a role in the inhibition of transcription and replication by VP24.

**Fig. 5.**
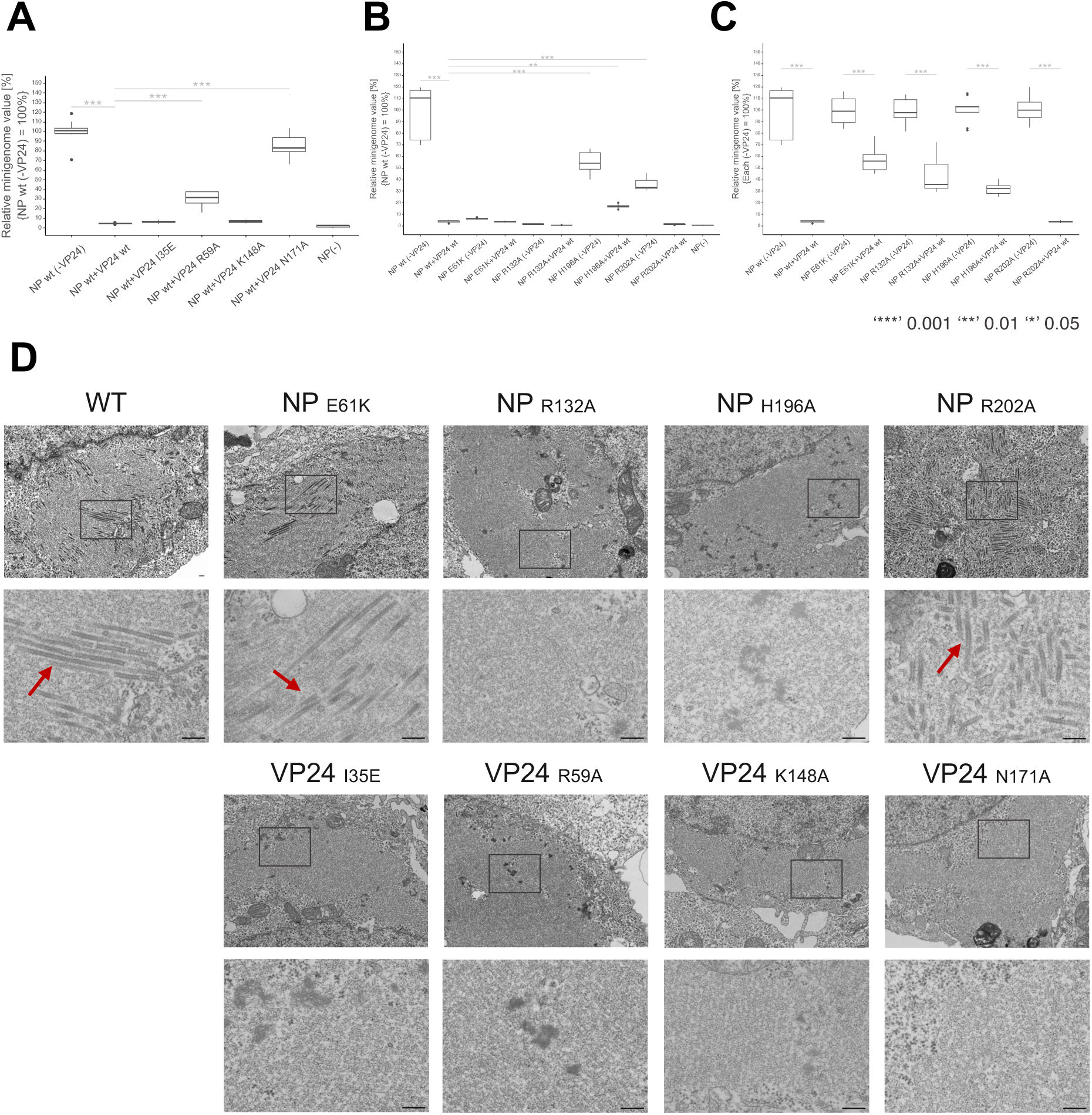
Effects of NP and VP24 mutations on RNA synthesis and NCLS formation. A-C. Transcription and replication activity evaluated using the minigenome assay with NP and VP24 mutants. The results are presented in box-and-whiskers plots. The plots detail the maxima, 75th percentile, median, 25th percentile, and minima, with outliers represented by dots. The activities are compared between (A) Wild-type VP24 and its mutants, with the activity of wild-type NP in the absence of VP24 normalized to 100%. (B) Wild-type NP and NP mutants, with the activity of wild-type NP normalized to 100%. (C) Wild-type NP and NP mutants, with the activity of each specific NP type in the absence of VP24 normalized to 100%. ***: *p* <0.001, **: *p* <0.01, *: *p* < 0.05 D. Ultrathin section electron microscopy of HEK293T cells co-expressing NP and VP24 mutants and VP35. Each lower panel shows a close-up view of the square area of the upper panel. Red arrows indicate the NCLS. Scale bars represent 100 nm.

To investigate whether each mutant is assembled into NCLS, we performed ultrathin section electron microscopy of cells co-expressing NP, VP24, and VP35. Similar to wild-type NP, NP E61K and NP R202A mutants formed tubular structures with dense walls approximately 50 nm in diameter, representing NCLS^3,4^ (Fig. 5D). As expected from the results obtained using IFA, VP24 N171A and R59A mutants did not form NCLS structures. However, other mutants, such as NP R132A and VP24 K148A (located at the interface between NP-2 and VP24-2) and VP24 I35E (located at the interface between VP24-1 and VP24-2), were also unable to form NCLS in cells (Fig. 5D). These results demonstrated that NCLS formation is not necessary for the inhibition of viral genome transcription and replication. Rather, it is likely that the interactions between VP24-1 and NP-1 via N171 and electrostatic interactions are important for its inhibitory activity. In addition, the interactions of VP24-2 with both NP-2 and VP24-1 are potentially important for NCLS formation, although the interaction between NP-1 and NP24-1 is probably a prerequisite.

### NCLS formation is necessary for its intracellular transport

NP, VP24, and VP35 are essential for the release of the nucleocapsid from IBs and the subsequent intracellular transport of the nucleocapsid^20,21,30^, which is a prerequisite for the packaging of the nucleocapsid into progeny virions mediated by the matrix protein VP40^4,31^. As NCLS formation was unrelated to the inhibition of viral RNA synthesis (Fig. 5A-C), we hypothesized that NCLS formation is required for its intracellular transport. Therefore, we co-expressed NP, VP35, and one of the VP24 mutants, along with VP30-GFP which was used to visualized nucleocapsid movement using immunofluorescence microscopy. Here, we focused on VP24 mutants that did not form the NCLS, yet showed different regulatory activities for transcription and replication: VP24 N171A, which did not inhibit transcription and replication; VP24 R59A, which showed reduced inhibitory activity for transcription and replication; and VP24 I35E, which inhibited transcription and replication. In the presence of wild-type VP24, the NCLSs exhibited long-distance intracellular movement (>5 µm) per cell, as reported previously^20^. In contrast, considerably fewer NCLSs exhibited long-distance movement in the presence of VP24, N171A, R59A, and I35E (Fig. 6A,B). Furthermore, signal trajectory analysis demonstrated that the NCLSs containing VP24 mutants exhibited significantly shorter movements than the nucleocapsid containing wild-type VP24 (Fig. 6C). These results suggested that intracellular NCLS transport is independent of the inhibition of vRNA transcription and replication; however, it relies on the formation of NCLS.

**Fig. 6.**
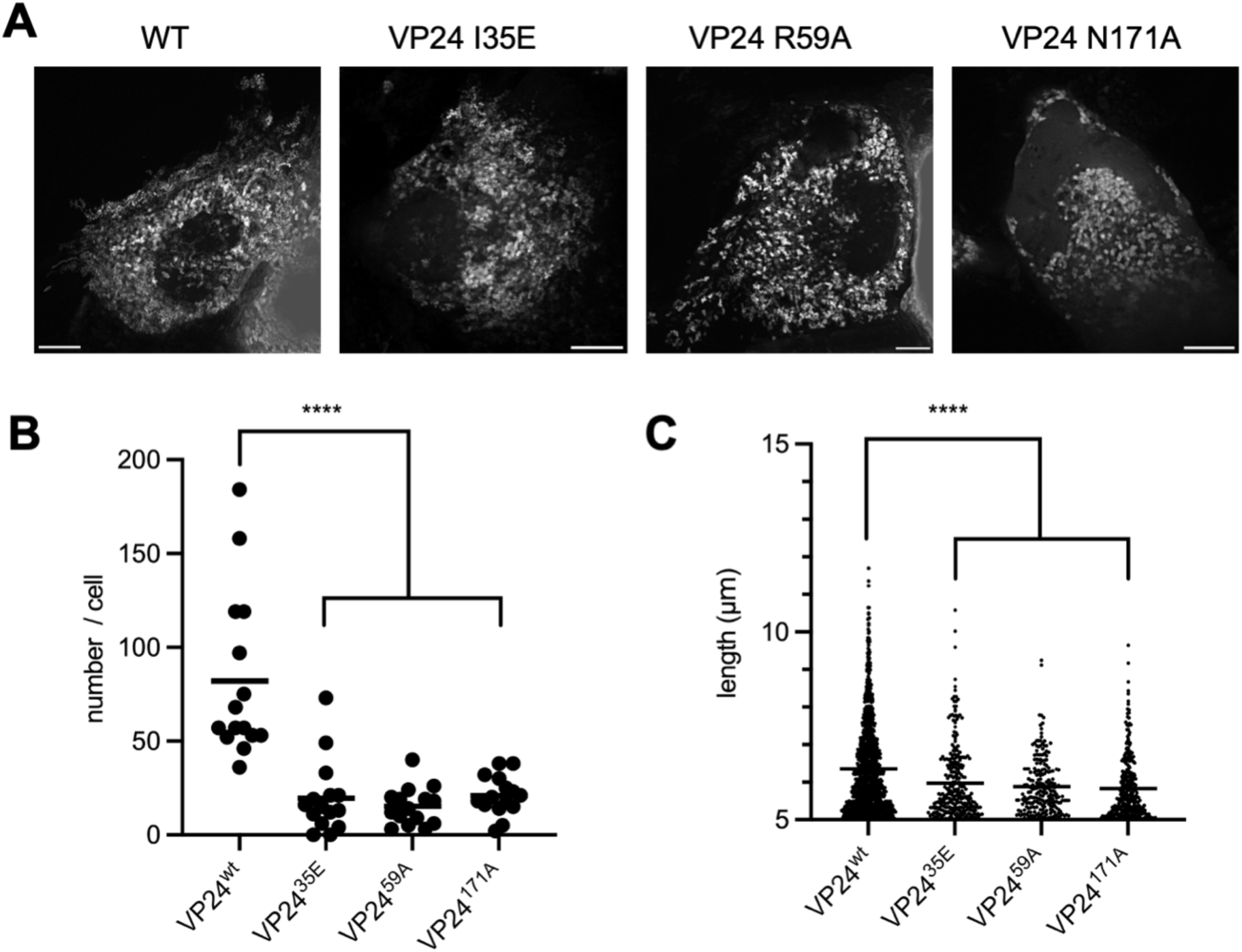
Intracellular transport of the NCLS composed of VP24 mutants. A. Maximum intensity projections of time-lapse images recorded for 3 min with 3 s intervals, showing the intracellular distribution of NCLS composed of wild-type VP24 and VP24 I35E, VP24 R59A, and VP24 N171A mutants. Scale bars represent 10 µm. B. Analysis of the number of NCLS showing long-range movement within the cell. Each point represents the number of moving NCLS per cell, with the average indicated by horizontal lines. ****: *p* <0.0001 C. Analysis of the trajectory length of NCLS movement within cells. Data points represent the length of each NCLS movement, with the trend showing a significant reduction in movement distance for VP24 mutants compared to wild-type. ****: *p* <0.0001

## Discussion

The EBOV nucleocapsid is essential for viral RNA transcription and replication, and genome packaging into progeny virions. Despite advances in understanding the respective structures of its components^8,14–17,22,26,27,32–35^ and the accumulated knowledge of nucleocapsid functions^12,13,16,19,21,23,25,33,36,37^, the structural basis of nucleocapsid functions remains poorly understood. Here, using cryo-EM single-particle analysis, we reported a 4.6 Å resolution cryo-EM structure of the EBOV NCLS within VLPs. Structure-based mutagenesis revealed specific interactions between the two NPs and two VP24s that govern the regulation of viral RNA synthesis, nucleocapsid assembly and intracellular nucleocapsid transport. This study deciphered the structure-function relationship of the nucleocapsid and revealed the distinct roles of each of the two VP24 molecules on the nucleocapsid during the viral life cycle, suggesting a more sophisticated regulatory mechanism than previously believed.

Based on our structural and functional analyses, we proposed a mechanistic model for the role of VP24s in regulating viral RNA synthesis and nucleocapsid formation (Fig. 7). Among the mutations located within the NP–VP24 interfaces, only the N171A mutation on VP24 eliminated the binding affinity of VP24 with NP (Fig. 4A), suggesting that the interaction between the 169-173 loop of VP24-1 and the C-terminal α-helix 18 of NP-1 is indispensable for NP–VP24 interactions. Thus, VP24 initially binds to NP-1 on the NP–RNA helix as VP24-1 via its 169-173 loop (Fig. 7A). Notably, VP24 R59A, similar to VP24 N171A (Fig. 4B), was not recruited into IBs, suggesting that VP24 R59 is likely important for stabilizing the NP-1–VP24-1 interaction following the initial NP-1–VP24-1 interaction via the 169-173 loop (Fig. 7A).

**Fig. 7.**
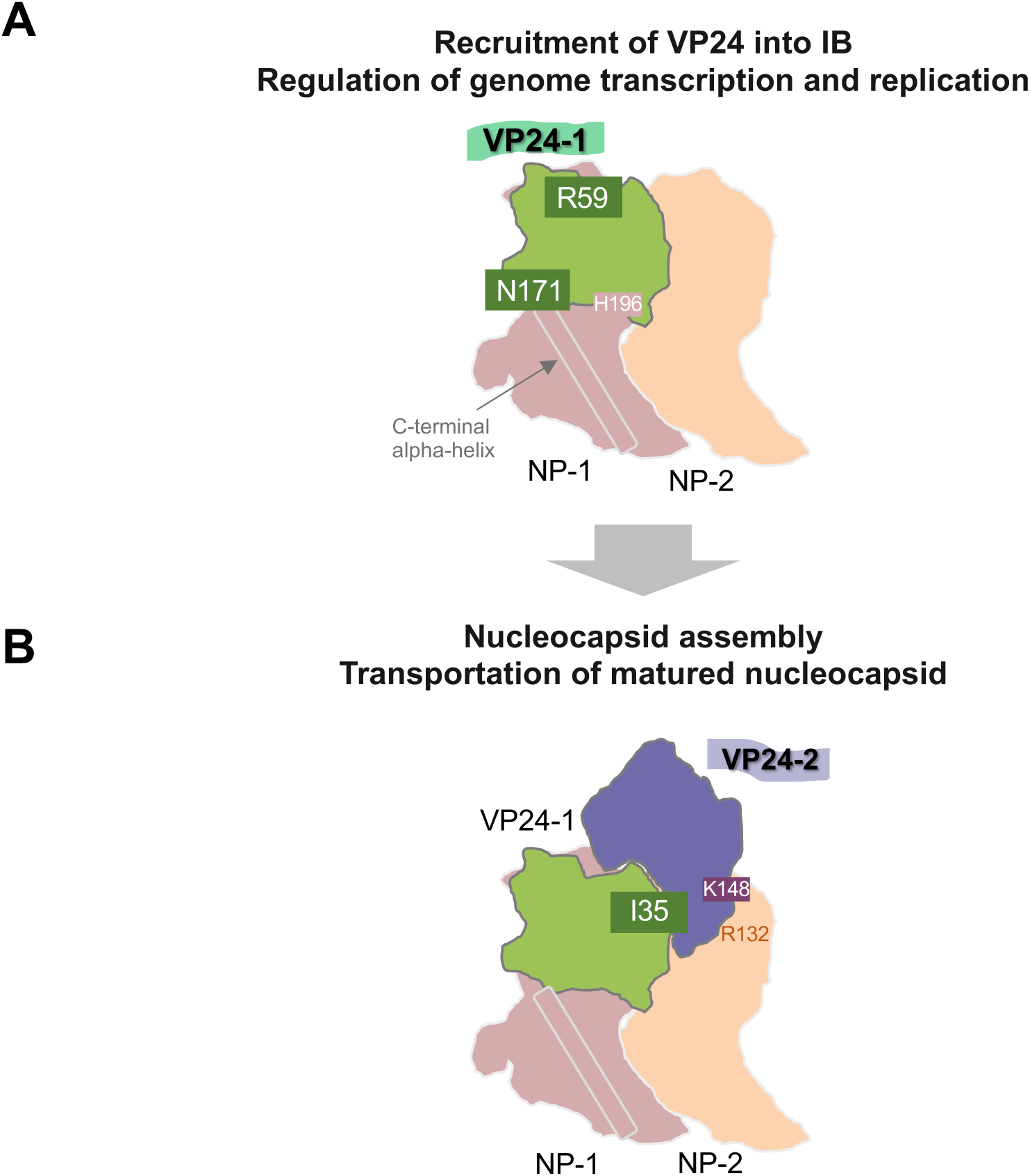
A proposed model of EBOV nucleocapsid assembly and the roles of two VP24 molecules. Our proposed model describes the two-phase strategy of NCLS formation and function. A. The first phase depicts the role of VP24-1 in initial nucleocapsid assembly, highlighting that the interface with NP-1 is critical for the recruitment of VP24 into inclusion bodies (IBs) and termination of genome transcription and replication. This diagram highlights the importance of residue N171 in VP24-1, as a critical binding site for NP. Moreover, VP24-1 R59 and potentially NP-1 H196 are involved in recruiting VP24 to IBs and inhibiting transcription and replication. B. The second phase illustrates the involvement of VP24-2 in the structural maturation of the nucleocapsid. Mutations in VP24 K148 and NP R132 at the NP-2–VP24-2 interface and VP24 I35 at the VP24-1–VP24-2 interface are crucial for nucleocapsid assembly. This stage highlights the necessity of these interactions for nucleocapsid formation and the subsequent transport of mature NCLS for the formation of progeny virions.

Using the minigenome assay, we found that VP24 (N171A and R59A) and NP (H196A and R202A), which have a mutation within the NP-1–VP24-1 interface, exhibited complete or partial failure to inhibit vRNA transcription and replication, whereas others that have a mutation within the NP-2–VP24-2 and the VP24-1–VP24-2 interface did not (Fig. 5A-C). These results strongly suggest that the binding of VP24-1 to NP-1 on the NP–RNA helix plays a critical role in inhibiting vRNA transcription and replication, rather than the previously believed mechanism involving NCLS formation (Fig. 7A). Although speculative, the proximity of VP24-1 to the template vRNA, closer than VP24-2 on the NP– RNA helix, suggested a potential steric hinderance on the movement of the L–VP35 polymerase complex for viral RNA synthesis. Further structural analyses of the nucleocapsid, including the L–VP35 complex, are required to understand the mechanisms underlying the inhibition of vRNA transcription and replication by VP24-1.

Our structural model demonstrates that VP24 N171, crucial for the NP–VP24 interaction, is not located within the NP-2–VP24-2 interface, suggesting that the interaction between VP24-2 and NP-2 is weaker than that between VP24-1 and NP-1. The initial binding of VP24-1 to NP-1 on the NP–RNA helix suggested that the binding of VP24-2 to NP-2 depends on the presence of VP24-1. It is anticipated to be stabilized by both electrostatic interaction with NP-2 and hydrophobic interaction with VP24-1. In our mutational analysis, VP24 K148A and NP R132A mutants, which have a mutation within the NP-2–VP24-2 interface, and the VP24 I35E mutant, which has a mutation within the VP24-1–VP24-2 interface, failed to form NCLS (Fig. 5D), suggesting the importance of VP24-2 for NCLS formation (Fig. 7B). Notably, VP24-1 is also crucial for NCLS formation, as mutants with VP24 and NP containing a mutation within the VP24-1–NP-1 interface were incapable of forming the NCLS. Such mutations (VP24 N171A, R59A, and potentially NP H196A) would hamper VP24 transport into IBs and the interaction between VP24-1 and VP24-2, leading to a failure in NCLS formation. Considering that the amino acids described in Fig. 7 are well conserved among filoviruses, especially between Ebola and Lloviu viruses (Extended Data Fig. 4A,B), the roles of each VP24 identified in this study are likely common among filoviruses.

Recently, Watanabe *et al.* reported the intracellular NCLS structure at 9 Å resolution using cryo-ET combined with sub-tomogram averaging, and suggested the existence of a C-terminal domain of VP35 on VP24-2 and an additional copy of the monomeric RNA-free NP on VP24-1^38^. Although similar additional maps were observed for both the VP24 protrusions in our cryo-EM map (Fig. 1C), we were unable to annotate these molecules owing to the flexibility of this region. The flexibility of this layer in our model may be due to differences in the environment between intracellular IBs and VLPs. This additional outer layer, where an additional copy of NP is likely involved in the intra-rung interactions of nucleocapsids, suggests additional structural and functional regulation of the nucleocapsid, which was not addressed in our study. Analyzing the interactions between RNA-free NP and VP35 in the additional layer, presumed to be involved in progeny virion assembly, will be a notable focus for future research.

In conclusion, we determined the *in situ* NCLS structure within the EBOV VLP and revealed the structural basis of their distinct functions during the EBOV life cycle. The identification of specific residues required for the respective nucleocapsid functions revealed potential targets for novel therapeutic interventions. Although our findings will advance this field, they also highlight the complexity of nucleocapsid assembly and functional regulation. This study opens avenues for further research, particularly to understand the dynamic interplay between nucleocapsid components and elucidating the mechanisms governing their multiple functions.

## Methods

### Cell culture

HEK293T (human embryonic kidney), HEK293, and Huh-7 cells were maintained in Dulbecco’s modified Eagle’s medium (D5796, Sigma-Aldrich, Burlington, MA, USA) supplemented with 10% fetal bovine serum (10270-106, Thermo Fisher Scientific, Waltham, MA, USA) at 37 °C and 5% CO_2_.

### Plasmids

Plasmids expressing EBOV proteins (pCAGGS-L, pCAGGS-GP, pCAGGS-VP24, pCAGGS-VP30, pCAGGS-VP35, pCAGGS-NP, and pCAGGS-NP-FLAG), the T7-driven EBOV minigenome encoding firefly luciferase (p3E5E-luc), T7 DNA-dependent RNA polymerase (pCAGGS-T7), and pTK *Renilla* luciferase were provided by Yoshihiro Kawaoka (University of Tokyo). pCAGGS-VP30-GFP for the VP30-GFP fusion protein was generated as previously described^20^.

Mutant EBOV protein-expressing plasmids were generated via site-directed PCR mutagenesis, and their sequences were confirmed using Sanger sequencing.

### Antibodies

The following antibodies were used for immunofluorescence microscopy: mouse Alexa Fluor 594-conjugated anti-FLAG-tag (M18-A59; MBL, Tokyo, Japan), rabbit anti-VP24 (provided by Hideki Ebihara at the National Institute of Infectious Diseases), and Alexa Fluor 488-conjugated anti-rabbit (A-11008; Thermo Fisher Scientific).

The following primary antibodies were used for western blot analysis (including samples from immunoprecipitation and minigenome assays): rabbit anti-EBOV NP (0301-012; IBT Bioservices, Rockville, MD, USA), rabbit anti-VP24 (provided by Yoshihiro Kawaoka), and mouse anti-beta-actin (ab8226; Abcam, Cambridge, UK).

The following secondary antibodies were used for western blot analysis (including samples from immunoprecipitation and minigenome assays): horseradish peroxidase (HRP)-conjugated anti-rabbit (NA934; Cytiva, Marlborough, MA, USA) and HRP-conjugated anti-mouse (NA931; Cytiva).

### Expression and purification of EBOV VLPs

HEK293T cells seeded in a 10 cm dish were transfected using TransIT-293 (Mirus Bio, Madison, WI, USA) with 670 ng pCAGGS-GP, 4 µg pCAGGS-NP, 1 µg pCAGGS-VP24, 1 µg pCAGGS-VP35, 4 µg pCAGGS-VP40, 4 µg p3E5E-luc, and 1.5 µg pCAGGS-T7. Three days post-transfection, VLPs in supernatants were fixed with 1% paraformaldehyde (PFA) and purified via ultracentrifugation using a 20% (w/v) sucrose cushion in Tris-HCl buffer (10 mM Tris-HCl (pH 8.0), 150 mM NaCl, 1 mM EDTA). The purified VLPs were resuspended in Tris-HCl buffer and diluted at 1.0 mg/ml.

### Cryo-electron microscopy: sample preparation and data collection

The total 2.5 µL sample was applied to both sides of glow-discharged 400-mesh R1.2/1.3 Cu grids (Quantifoil, Jena, Germany). The grids were blotted for 14 s at a humidity of 100% at 8 °C and rapidly frozen in liquid ethane using a Vitrobot Mark IV (Thermo Fisher Scientific). Cryo-EM samples were initially screened using a Glacios cryo-EM operated at 200 kV and equipped with a Falcon4 camera installed at the Institute for Life and Medical Sciences, Kyoto University. Cryo-EM micrographs were acquired using a Titan Krios cryo-EM (Thermo Fisher Scientific) operated at 300 kV and equipped with a Gatan BioContinuum energy filter and Cs corrector (CEOS GmbH, Heidelberg, Germany) installed at the Institute for Protein Research, Osaka University. Data were collected using the Serial-EM software^40^ (v4.0) with a beam-image shift imaging scheme, with a K3 detector in counting and correlated double sampling imaging mode. The collection was performed at a magnification of 81,000 × and a pixel size of 0.88 Å. The slit width of the energy filter was set to 20 eV. Each micrograph was exposed to a total dose of 60 e-per Å², fractionated into 60 frames. The nominal defocus range for the dataset was –0.6 to –2 μm. The detailed imaging conditions are described in Extended Data Table 2.

### Cryo-electron microscopy: image processing

The detailed workflow is shown in Extended Data Fig. 2. The frames of each movie were aligned within RELION 4.0-beta^41,42^ on 5 × 5 patches, and contrast transfer function (CTF) estimations were performed using CTFFIND4^43^. The coordinates of the NCLS were manually selected from 5,897 micrographs. In total, 263,789 segments were extracted using a 720 × 720 pixel box size and an inter-box distance of 10%. The segments underwent two rounds of 2D classification in RELION 4.0-beta. The initial 3D model was generated from a previously reported cryo-ET map, EMD3871, employing a 60 Å low-pass filter. Following a single round of 3D classification, without a local search for helical symmetry, 34,865 segments (binned at 1.2375 Å/pix) underwent further 3D refinement. Additional CTF refinement, Bayesian polishing, and additional 3D refinement, incorporating a local search for symmetry and SIDESPLITTER^44^, yielded a 7.5 Å cryo-EM map. Helical parameters were locally searched based on previously reported values^8^, resulting in a –28.1059° twist and a 5.8984 Å rise.

To enhance the structure of the NCLS repeating unit, we expanded the helical symmetry and subtracted signals outside the NCLS from the particles for subsequent classification and local refinement. Using the relion_particle_symmetry_expand command, symmetrized helical segments were expanded into each repeating unit (418,380 particles) according to their helical symmetry, ensuring an interbox distance to prevent duplicates in the new particle sets. Subsequently, the signal outside the repeating unit (approximately 4-mer NPs + protrusions) was subtracted with a box size of 256 × 256 pixels. Thereafter, 3D classification without alignment was performed, focusing specifically on the repeating unit region with a mask. Finally. In total, 204,348 particles were imported into cryoSPARC^45,46^ (v3.3.1) for local refinement. The resulting cryo-EM map had an overall resolution of 4.6 Å.

### Structural model building and refinement

For the initial model, we docked previously reported structures, PDB ID: 5Z9W for NP and PDB ID: 4M0Q for VP24, onto the main-chain scaffold of PDB ID: 6EHM. To precisely refine the atomic models in the intermolecular region, the two molecules surrounding the NPs were additionally fitted to our cryo-EM map. The initial refinement process was executed via Rosetta using Relax^47–49^. A total of 100 models were generated, and those with better energy scores and fit densities were manually inspected. Small modeling errors were manually corrected in COOT (v0.9.8.7)^50,51^, and in the refinement was reiterated in Rosetta using Relax. The final model was refined using Phenix^52^ with phenix.real_space_refine, followed by manual correction using COOT and ISOLDE (v1.6.0)^53^. After removing the unmodeled region, model validity was assessed using Phenix (v1.20) and MolProbity^54^ software. Detailed results of the atomic modeling and evaluation are shown in the Extended Data Table. 2. The cryo-EM map and atomic models were displayed using UCSF Chimera X^55^ (v1.5) software. A contact map (Fig. 3A) was generated using the MAPIYA webserver^56^.

### Protein immunoprecipitation

HEK293T cells were seeded in 6-well plates and transfected with FLAG-tagged NP and/or VP24 expression plasmids (1 µg/well each) using TransIT-293. Two days post-transfection, cells were washed with cold phosphate-buffered saline (PBS). Cells were resuspended in 400 μL of lysis buffer (50 mM Tris-HCl [pH 8.0], 150 mM NaCl, 1% NP-40, cOmplete EDTA-free protease inhibitor tablets [Roche, Basel, Switzerland]), incubated for 1 h at 4 °C, and centrifuged at 12,000 × *g* for 10 min at 4 °C. The supernatant was incubated on a rotator overnight at 4 °C with an additional 16 μL of anti-FLAG M2 magnetic beads (Sigma-Aldrich. The beads were washed six times with wash buffer (50 mM Tris-HCl [pH 8.0], 150 mM NaCl, cOmplete EDTA-free protease inhibitor tablets) and subsequently eluted in 150 μL of wash buffer with 500 ng/μL FLAG peptide (Merck, Darmstadt, Germany) by rotation on a rotator for 2 h at 4 °C. The cell lysates and eluates were electrophoresed on a sodium dodecyl sulfate-polyacrylamide gel and subjected to western blot analysis.

### Minigenome assay

HEK293T cells were cultured in 24-well plates. Each well was transfected with plasmids encoding EBOV nucleocapsid components, including 1,000 ng pCAGGS-L, 75 ng pCAGGS-VP30, 100 ng pCAGGS-VP35, 100 ng pCAGGS-NP, 200 ng firefly luciferase p3E5E-luc, 200 ng T7 DNA-dependent RNA polymerase (pCAGGS-T7), and 10 ng *Renilla* luciferase (pTK r.luc), with or without 100 ng pCAGGS-VP24. Transfection was performed using 4 µL TransIT 293 (Takara, Shiga, Japan) and 100 µL OPTI-MEM (Thermo Fisher Scientific). At 48 h post-transfection, cells were lysed using Passive Lysis buffer (Promega Madison, WI, USA). Luciferase activities were measured using a GloMax®-Multi+ Detection System (Promega), along with Luciferase Assay Reagent II and Stop & Glo® Reagent (Promega), following the manufacturer’s protocol.

### Ultrathin section electron microscopy

HEK293T cells were seeded on a 6-well plate, transfected with 5 µg NP or NP mutant-encoding plasmids, 1 µg VP24 or VP24 mutant-encoding plasmids, and 1 µg VP35-encoding plasmids. At 24 h post-transfection, cells were fixed with aldehydes, postfixed with 1 % osmium tetroxide, dehydrated in a graded ethanol series, and embedded in an EPON 812 (TAAB, Berks, UK). Ultrathin sections were stained with uranyl acetate and lead citrate and observed using a Hitachi HT-7700 microscope operated at 80 kV (Hitachi Hi-Tech, Tokyo, Japan).

### Immunofluorescence assay

HEK293 cells were plated in a 35 mm glass-bottom dish (Matsunami, Osaka, Japan) coated with collagen I (Corning, Corning, NY, USA). Cells were transfected with 1 µg/each of FLAG-tagged NP, VP24, and VP35 expressing plasmids. At 48 h post-transfect, transfected cells were fixed in 4% PFA in PBS for 15 min and subsequently permeabilized with 0.1% Triton X-100 (MP Biomedicals, Santa Ana, CA, USA) in PBS for 15 min. Cells were blocked with Blocking One solution (Nacalai Tesque, Kyoto, Japan) for 30 min, followed by overnight incubation at 4 °C with primary antibodies [anti-FLAG-tag-Alexa Fluor 594 (MBL, Tokyo, Japan) 1:1000; rabbit anti-EBOV-VP24^21^ 1:400]. Thereafter, cells were incubated with secondary antibodies [Alexa Fluor 488 anti-rabbit (Thermo Fisher Scientific) 1:1000] for 1 h at room temperature. Cells were treated with Hoechst 33342 (1:1000; Thermo Fisher Scientific) for nuclei staining following the application of secondary antibodies. Section images were recorded using DeltaVision Elite (Cytiva, Marlborough, MA, USA) with a 60× objective and subsequently deconvolved and projected using SoftWoRx Version 6.5.2.

### Live-cell microscopy

Huh-7 cells (1 × 10^4^ cells) were seeded onto an 8-well slide (Ibidi, Gräfelfing, Germany). Transfection with plasmids expressing VP30-GFP, NP, VP24 or VP24 mutant, and VP35 was performed in 50 μL Opti-MEM (Thermo Fisher Scientific). The inoculum was removed at 1 h post-transfection, and a volume of 250 μL CO_2_-independent Leibovitz’s medium (Thermo Fisher Scientific) was added. Live-cell imaging was performed using a Keyence BZ-X 810 (KEYENCE, Osaka, Japan) with a 100× oil objective lens at 20 h post-transfection. Pictures and movie sequences were processed using Fiji software^57^. For trajectory analysis, 15 cells per sample were analyzed using Imaris 10.0.0 software (Oxford Instruments, Abingdon, UK). To prevent contamination of signals exclusively from VP30, we omitted signals demonstrating a trajectory length <5 μm, as they deviate from the typical transport pattern of NCLSs^20,58^.

### Statistical analyses

To assess statistically significant differences in the minigenome assay, we employed ANOVA-Dunnett’s test to correct for multiple hypotheses using the RStudio software. The results are reported as the mean ± s.d. with *p* <0.01 considered statistically significant. For the NCLS trajectory analysis of live-cell imaging, statistical analysis was performed using the Student’s t-test with GraphPad Prism version 10.0.0 for MacOS (GraphPad Software, Boston, MA, USA, www.graphpad.com), and *p* <0.0001 was considered statistically significant (****). All experiments were performed in triplicate.

## Author’s contributions

Y.F-F., Y.S., and T.N. designed the study. Y.F-F., S.H., A.H., Y.T., N.Y., K.H., and Y.S. performed the experiments. Y.F-F., S.H., Y.T., and Y.S. analyzed the data. Y.F-F., S.H., Y.S., and T.N. wrote the manuscript. All the authors have reviewed and approved the final manuscript.

## Data availability

The data in this study are available from the corresponding author upon request. The cryo-EM maps generated in this study have been deposited in the Electron Microscopy Data Bank under accession number EMD-XXXXX. The atomic coordinates reported in this paper have been deposited in the Protein Data Bank (PDB) under accession number XXXX. Raw movies were deposited in the Electron Microscopy Public Image Archive under the accession number EMPIAR-XXXXX.

## Acknowledgements

We thank Hideki Ebihara (National Institute of Infectious Diseases) and Satoko Yamaoka (Mayo Clinic) for providing the VP24 antibody and Yoshihiro Kawaoka (University of Tokyo) for providing plasmids expressing EBOV viral proteins and the minigenome. This work was supported by a Grant-in-Aid from the Japan Society for the Promotion of Science (JSPS) Fellows (21J12207) and an ANRI fellowship (to Y.F-F.), the Otsuka Toshimi Scholarship (to S.H.); Grant-in-Aid for the Ministry of Education, Culture, Sports, Science and Technology (MEXT) Leading Initiative for Excellent Young Researchers, JSPS Scientific Research (C) (21K07052), Japan Science and Technology Agency (JST) FOREST Program (JPMJFR214S) (to Y.S.); Japan Agency of Medical Research and Development (AMED: JP23fm0208101, JP22fm0208101, 22wm0325023h9903), JSPS (22KK0115, 21K07059), MSD Life Science Foundation, Naito Foundation, Takeda Science Foundation (to Y.T.); and JSPS Core-to-Core Program A, Advanced Research Networks (JPJSCCA20190008), JSPS Grant-in-Aid for Challenging Exploratory Research (22K19431), JSPS Grants-in-Aid for Scientific Research, Fund for the promotion of Joint International Research (22KK0115), JST Core Research for Evolutional Science and Technology (JPMJCR20HA), AMED (JP23fm0208101 and 22fk0108552h0001), the cooperative research project program of the National Research Center for the Control and Prevention of Infectious Diseases, Nagasaki University (to T.N.); Grant for Joint Research Project of the Institute of Medical Science at the University of Tokyo, Joint Usage/Research Center Program of the Institute for Life and Medical Sciences at Kyoto University, the Takeda Science Foundation (to Y.S. and T.N.). This study was also supported by the Platform Project for Supporting Drug Discovery and Life Science Research (Basis for Supporting Innovative Drug Discovery and Life Science Research (BINDS)) from AMED under grant number JP23ama121001 (support number 5889).

