## Extended Figures and Tables for "Structural basis of Ebola virus nucleocapsid assembly and functions"

### Extended Figures and legends

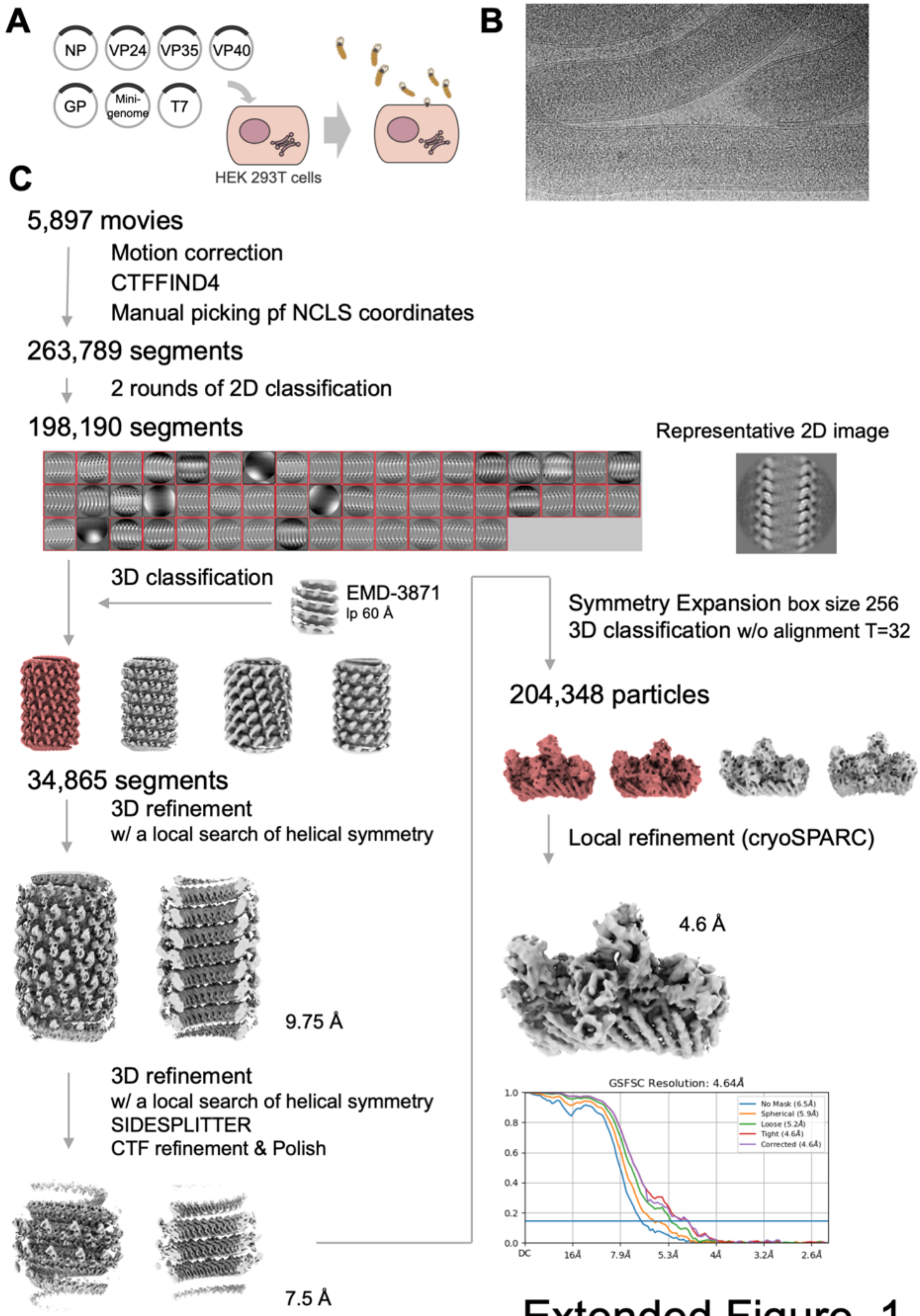

Extended Figure. 1

**Extended Data Fig. 1 Cryo-EM data processing of the NCLSs in VLPs.**

A. Schematic representation of the VLPs production in HEK 293T cells.

B. Representative cryo-EM image of purified VLPs.

C. Summary of the cryo-EM data processing workflow. Most processes executed using RELION software, except for local refinement with cryoSPARC.

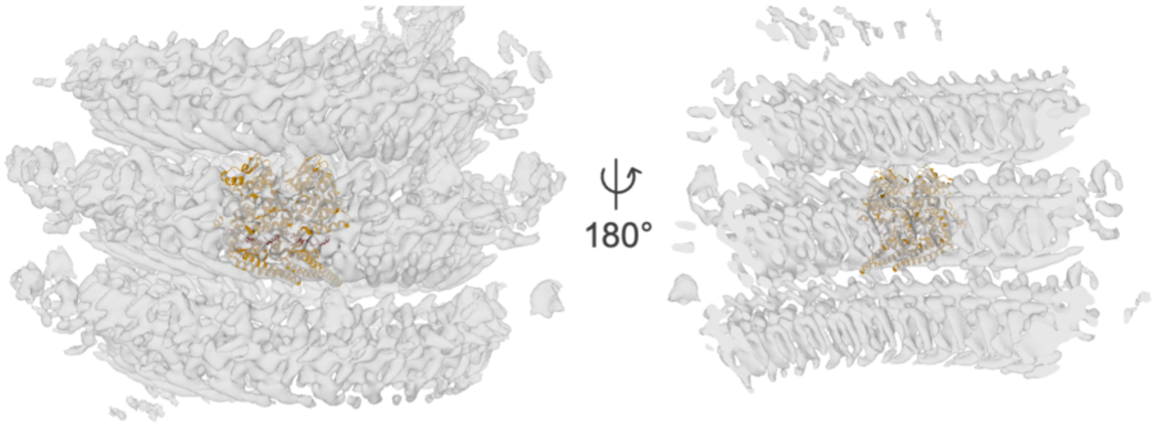

Extended Figure. 2

**Extended Data Fig. 2 Comparative visualization of the EBOV NCLS obtained via cryo-EM and the previously determined NP–RNA complex.**

The cryo-EM map obtained in this study using helical symmetry (shown in gray) is superimposed on the atomic model of the purified NP–RNA complex structure[22] (PDB ID: 5Z9W) shown in orange. Density thresholds were set to indicate the inner helical densities.

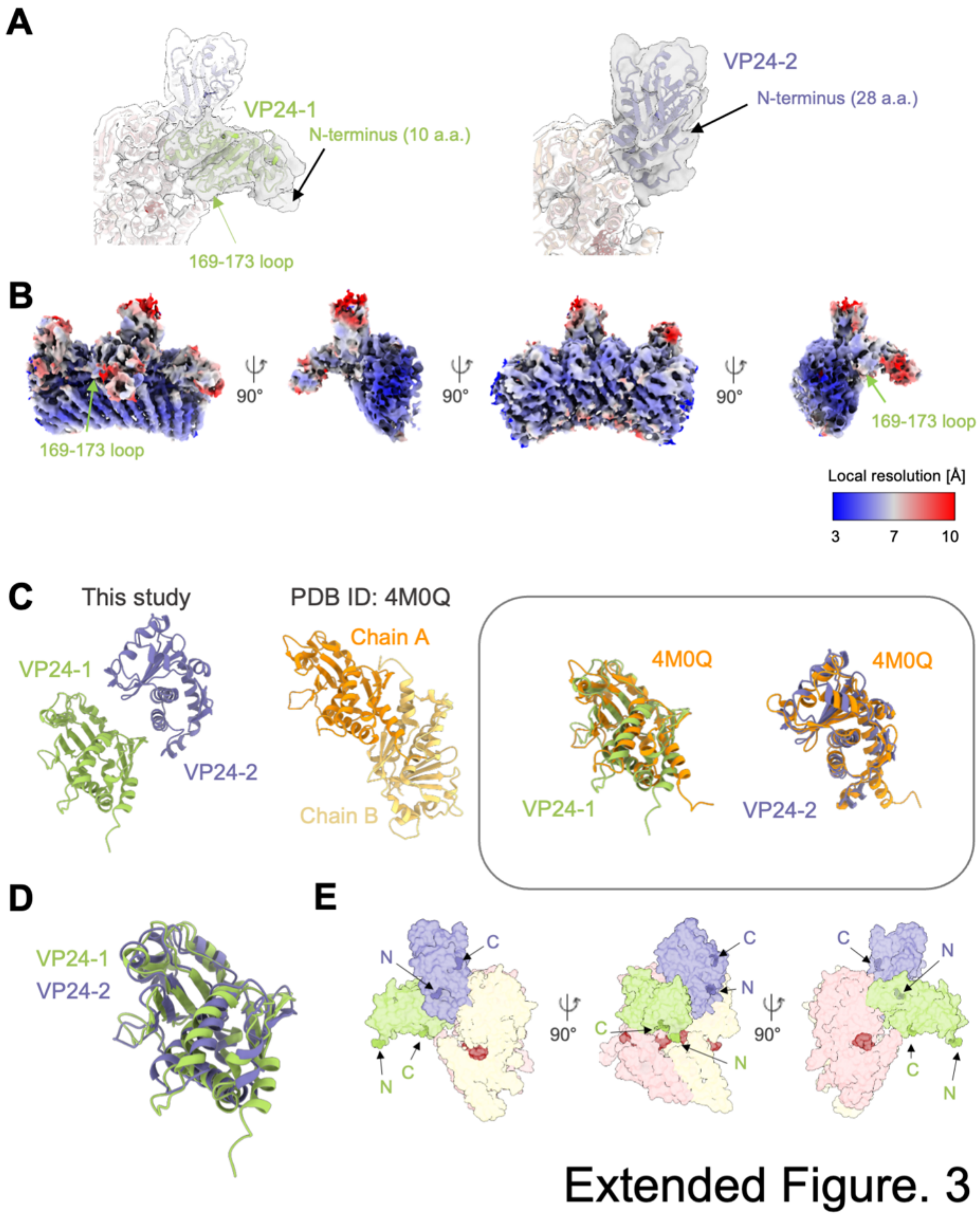

**Extended Data Fig. 3 Detailed structural comparison of VP24 interaction interfaces, comparing them with previously reported structures**

A. Close-up views of VP24-1 (10 a.a. to 231 a.a.) and VP24-2 (28 a.a. to 231 a.a.), with cryo-EM map (shown in gray). A black arrow indicates each N-terminus, and a light green arrow indicates the 169-173 loop.

B. Local resolution map of the helical repeating unit. A light green arrow indicates the 169-173 loops. The surface is colored according to the local resolution as defined by the color map to the right.

C. Comparison of our VP24-1 and VP24-2 models from this study with the crystal structure of EBOV VP24 in the dimer form[26] (PDB ID: 4M0Q). The right panel shows the structural comparisons of our VP24 models from the PDB 4M0Q chain A.

D. Structural comparison of VP24-1 and VP24-2 from this study.

E. Surface representation with highlighted N- and C-termini of VP24s.

A

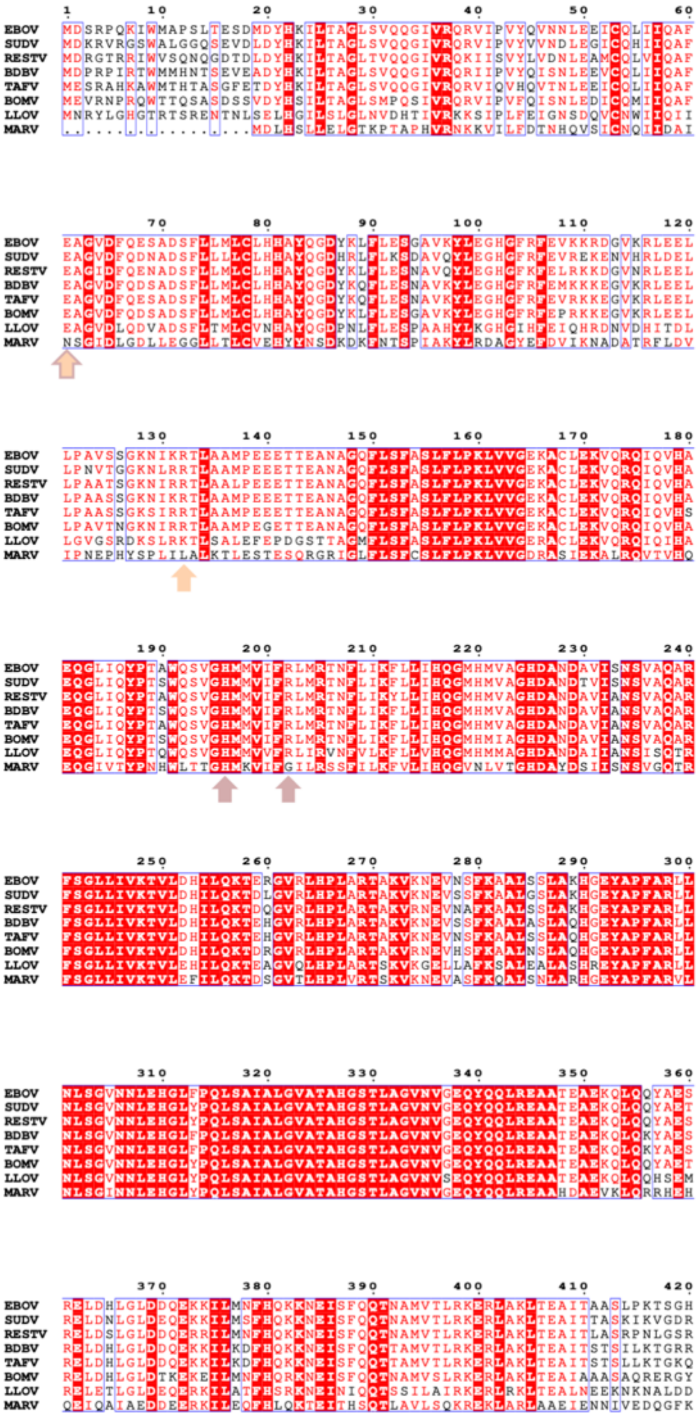

**B**

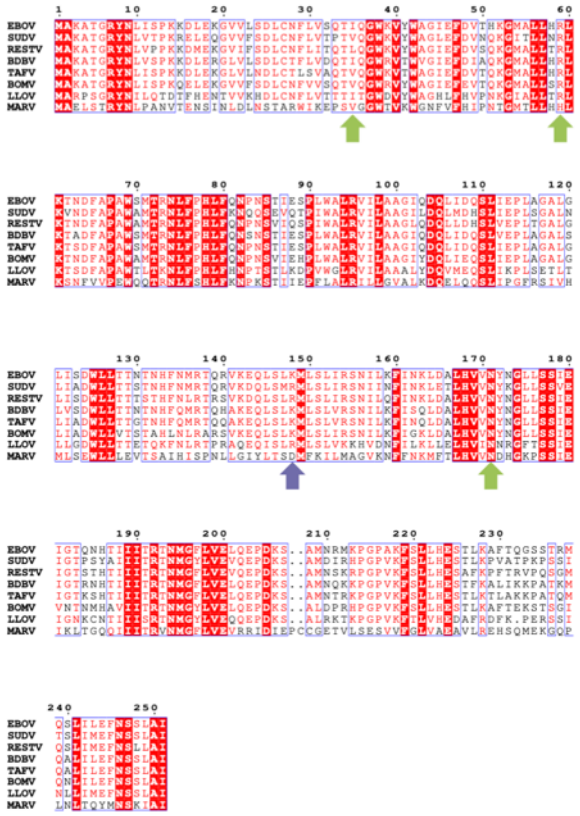

Extended Figure. 4

**Extended Data Fig. 4 Sequence alignment of the NP and VP24 among filoviruses**

The amino acid sequences of filovirus (A) NP and (B) VP24 (EBOV: *Zaire ebolavirus* {NCBI accession number: P18272 and Q05322}, SUDV: *Sudan ebolavirus* {NCBI accession number: Q9QP77 and Q5XX02}, RESTV: *Reston ebolavirus* {NCBI accession number: Q8JPY1 and Q77DB4}, BDBV: *Bundibugyo ebolavirus* {NCBI accession number: B8XCM7 and B8XCN4}, TAFV: *Taï Forest ebolavirus* {NCBI accession number: B8XCN6 and B8XCP3}, BOMV: *Bombali ebolavirus* {NCBI accession number: A0A4D5SFX8 and A0A343EQF5}, LLOV: *Lloviu cuevavirus* {NCBI accession number: G8EFI1 and G8EFI8}, MARV: *Marburg marburgvirus* {NCBI accession number: Q1PD53 and Q1PD62}) were aligned with ClustalW2.1[59] and visualized using ESPript 3.0[60]. The number of alignments was assigned based on EBOV sequences. Conserved residues are highlighted in red, similar residues are colored in red, and both residues are outlined in blue. The amino acids mentioned in the manuscript are indicated by arrows in the same color, as shown in Fig. 1.

**Extended Table 1. Comparison of helical parameters in EBOV NC-related structures**

| | This study | EMD-6903 | EMD-3871 | EMD-3873 |
|---|---|---|---|---|
| Target | VLP | NP–RNA complex | VLP | Infectious virion |
| Method | Single-particle analysis | Single-particle analysis | cryo-ET | cryo-ET |
| Pitch | 75.55 Å | 73.56 Å | 75 ± 3 Å | 74 ± 1 Å |
| Number of unit per turn | 12.8 | 24.4 for 1-mer NP means 12.2 for 2-mer NPs | 12.8 or 13.8 | 11.9 or 12.9 |

**Extended Table 2. Cryo-EM data collection, refinement and validation statistics**

| | EBOV NCLS in VLPs (EMDB-XXXXX) (PDB ID XXXX) |
|---|---|
| **Data collection and processing** | |
| Voltage (kV) | 300 |
| Electron exposure (e–/Å$^2$) | 60 |
| Defocus range (μm) | -0.625 to -2 |
| Pixel size (Å) | 0.88 |
| Initial particle images (no.) | 263,789 |
| Final particle images for the helical reconstruction (no.) | 34,865 |
| Final particle images for the symmetry expanded particles (no.) | 204,384 |
| Map resolution for the symmetry expanded particles (Å) | 4.6 |
| FSC threshold | 0.143 |
| **Refinement** | |
| Initial model used (PDB code) | 5Z9W, 4M0Q, 6EHM |
| Model resolution (Å) | 4.5 |
| FSC threshold | 0.143 |
| Model resolution range (Å) | 4.2-6.9 |

| | |
|---|---|
| Map *B* factor | 173.9 |
| Model composition | |
| Non-hydrogen atoms | 9776 |
| Protein residues | 1206 |
| Nucleotide | 12 |
| *B* factors (Å$^2$) | |
| Protein | 264.16 |
| Nucleotide | 225.73 |
| R.m.s. deviations | |
| Bond lengths (Å) | 0.012 |
| Bond angles (°) | 2.117 |
| Validation | |
| MolProbity score | 1.49 |
| Clashscore | 0.92 |
| Poor rotamers (%) | 1.75 |
| Ramachandran plot | |
| Favored (%) | 90.65 |
| Allowed (%) | 7.43 |
| Disallowed (%) | 1.92 |
